# Intrinsic neuronal activity during migration controls the recruitment of specific interneuron subtypes in the postnatal mouse olfactory bulb

**DOI:** 10.1101/2020.07.28.224568

**Authors:** Bugeon Stéphane, Haubold Clara, Ryzynski Alexandre, Cremer Harold, Platel Jean-Claude

## Abstract

Neuronal activity has been identified as a key regulator of neuronal network development, but the impact of activity on migration and terminal positioning of interneuron subtypes is poorly understood. The absence of early subpopulation markers and the presence of intermingled migratory and post-migratory neurons makes the developing cerebral cortex a difficult model to answer these questions. Postnatal neurogenesis in the subventricular zone offers a more accessible and compartmentalized model. Neural stem cells regionalized along the border of the lateral ventricle produce two main subtypes of neural progenitors, granule cells and periglomerular neurons that migrate tangentially in the rostral migratory stream before migrating radially in the OB layers. Here we take advantage of targeted postnatal electroporation to compare the migration of these two population. We do not observe any obvious differences regarding the mode of tangential or radial migration between these two subtypes. However, we find a very striking increase of intrinsic calcium activity only in granule cell precursors when they switch from tangential to radial migration. By decreasing neuronal excitability in granule cell precursors, we find that neuronal activity is critical for normal migratory speed at the end of tangential migration. Importantly, we also find that activity is required for normal positioning and survival of granule cell precursors in the OB layers. Strikingly, decreasing activity of periglomerular neuron precursors did not impact their positioning or survival. Altogether these findings suggest that neuronal excitability plays a subtype specific role during the late stage of migration of postnatally born olfactory bulb interneurons.

**Significance Statement:** While neuronal activity is a critical factor regulating different aspects of neurogenesis, it has been challenging to study its role during the migration of different neuronal subpopulations. Here, we use postnatal targeted electroporation to label and manipulate the two main olfactory bulb interneuron subpopulations during their migration: granule cell and periglomerular neuron precursors. We find a very striking increase of calcium activity only in granule cell precursors when they switch from tangential to radial migration. Interestingly, blocking activity in granule cell precursors affected their migration, positioning and survival while periglomerular neuron precursors are not affected. These results suggest that neuronal activity is required specifically for the recruitment of granule cell precursors in the olfactory bulb layers.

## Introduction

Neuronal networks comprise different subtypes of interconnected neurons in a precise region-specific stoichiometry and organization. For example, in the mouse neocortex about 85% of all neurons are glutamatergic, whereas 15% are inhibitory GABAergic interneurons. The mechanisms that regulate the migration and recruitment of appropriate numbers of inhibitory neurons in precise brain regions remain largely unknown.

Electrical activity has been identified as a key regulator of network development. For example, activity controls numerical population matching in the cortex. Electrical input from pyramidal neurons to interneurons determines their survival or death thereby controlling the amount of interneurons to the needs of the principal neuron pool (Lodato et al., 2011; Wong et al., 2018). In addition, there are evidence that electrical activity impacts on neuronal migration and consequently final positioning of cortical interneurons. For example, specific subpopulations of interneurons show a shift in laminar positioning when their activity is inhibited by the expression of the Kir2.1 channel (De Marco Garcia, 2011).

However, despite such first observations, the specific role of neuronal activity during migration is not well understood. This is mainly due to a paucity of early markers, that allow the identification of interneuron subpopulations already during their migratory phase and before reaching terminal positions. Moreover, specifically during corticogenesis migratory and post-migratory populations are intermingled and homogeneous compartments are not identifiable. Thus, to understand the impact of activity on interneuron migration and terminal positioning, lineage tracing and developmental models with clear compartmentalization are needed.

Activity has also been shown to be a major regulation factor of interneuron integration in the olfactory system. Indeed, in the postnatal and adult rodent, new interneurons are produced throughout life (Lois and Alvarez-Buylla, 1994) and permanently added to the preexisting circuitry of the olfactory bulb (OB) (Platel et al., 2019). In this system neural stem cells localized in the V-SVZ along lateral ventricles generate neuronal precursors that migrate tangentially via the rostral migratory stream (RMS) into the core of the OB. Here they detach from the RMS and switch from tangential to radial migration. Granule Cell Precursors (GC-P) migrate short distances and settle in the Granule Cell Layer (GCL). These represent approximately 90% of all new neurons. Periglomerular Neuron Precursors (PGN-P) traverse the entire GCL to reach the peripheral Glomerular Layer (GL) before their integration. Importantly, the generation of these interneuron subtypes relies on the regionalization of the stem cell compartment in the V-SVZ. For example, GCs are lineage related to stem cells localized in lateral aspects of the ventricular wall, whereas dopaminergic/GABAergic, calretinin-positive/GABAergic and pure Gabaergic PGN are predominantly derived from stem cells in the dorsal and septal walls (Merkle et al., 2007; Fernandez et al., 2011). Several lines of evidence suggest that, like in the cortex, activity is a major regulator of this interneuron integration process. For example, increasing neuronal activity via olfactory training (Mouret et al., 2008) or by over-expression of the bacterial sodium channel NaChBac expression (Lin et al., 2010) enhance the number of newborn neurons integrated. By contrast, sensory deprivation via naris occlusion decreased the survival of newborn neurons in the OB (Saghatelyan et al., 2005; Yamaguchi and Mori, 2005; Platel et al., 2019).

Altogether, the highly compartmentalized environment of the OB neurogenic system provides the possibility to trace interneuron subpopulations destined for different target layers and study their specific migratory behavior and activity patterns during tangential and radial phases. Finally, due to the fact that defined stem cell pools can be efficiently manipulated, it is particularly well suited to analyze and manipulate activity patterns.

Here we use targeted brain electroporation to label specifically the lateral and septal ventricular wall and show that this approach labels with high predominance either future GCs or PGN. We compared their migratory behavior and neuronal activity at high resolution, combining acute forebrain slices with 2-photon microscopy. Tangential and radial migration was indistinguishable between both populations. However, GC-P showed increased calcium activity already in the distal RMS, before exit and onset of radial migration. Moreover, we find that inhibition of activity in GC-P but not in PGN-P affects the transition from tangential to radial migration, their final positioning and survival. Our results indicate that activity during migration is specifically implicated in the recruitment process of GC-P before they enter the OB neuronal network. In constrast, the migration of PGN-P to the OB network seems largely independent on activity.

## Results

### A new model to study interneuron migration

Previous work showed that terminal positioning of interneurons in the OB is dependent on the position of their respective stem cells along the walls of the lateral ventricles. Neurons born along the lateral wall tend to remain in the deep positioned GCL whereas neurons derived from the medial wall preferentially migrate into the peripheral GL (Merkle et al., 2007; Fernandez et al., 2011).

In order to quantify this correlation, we used targeted in vivo brain electroporation, which allows specific introduction of genetic material in the lateral or medial stem cell compartments (Fernandez et al., 2011; Bugeon et al., 2017). Animals were electroporated at P1 and the OB was analysed 21 days later (21 days post electroporation, dpe) when newborn neurons reached the OB and settled in their terminal positions. For clarity we use false-colour representations, lateral electroporation in green and medial in red, throughout the manuscript. Introduction of a GFP-expression plasmid pCAG-GFP into the lateral ventricular wall labelled with high predominance (about 80%) granule cells that remained in central positions within the GCL. Only few labelled neurons were found in the MCL and the GL (Fig. 1A, C). In contrast, electroporation of pCAG-GFP into the medial wall labelled mainly cells that traversed the GCL to settle in the GL as PGN (about 80%; Fig. 1B,C).

**Figure 1:**
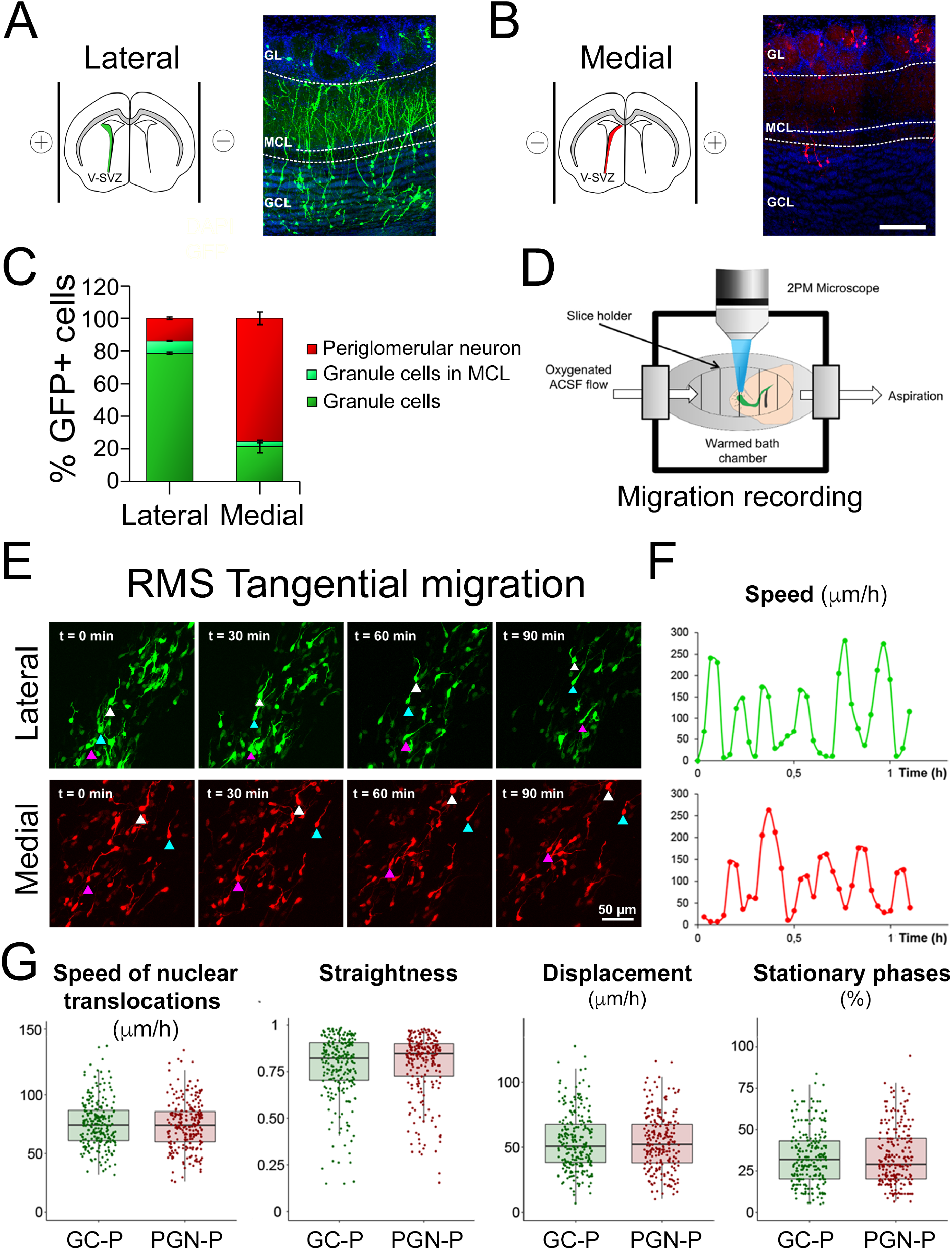
Tangential migration recording of postnatally generated subpopulation of olfactory bulb interneurons. **A.** Left, Schematic of the V-SVZ from animals electroporated laterally. Right, coronal sections of the OB at 21 dpe after lateral electroporation at P1 with a GFP encoding plasmid. Most of the neurons generated by lateral stem cells are located in the granule cell layer. **B.** Left, Schematic of the V-SVZ from animals electroporated medially. Right, coronal sections of the OB at 21 dpe after medial electroporation at P1 with a GFP encoding plasmid. Most of the neurons generated by medial stem cells are located in the glomerular layer. Scale bar: 50 μm. **C.** Quantification of the mean percentage of GFP positive cells in the different OB layers after lateral or medial electroporation. **D.** Experimental setup for 2-photon time lapse imaging of acute brain slices. **E.** Snapshots of time-lapse two-photon imaging of lateral and medial cells respectively. For each panel, arrowheads indicate individual cells followed over time. Time is indicated in minutes. Scale bar: 50 μm. **F.** Example of instantaneous speed recorded during a 1-hour movie for a lateral and a medial cell. **G.** Mean speed of nuclear translocations, straightness, displacement per hour and percentage of stationary phases for RMS tangentially migrating lateral and medial cells. n = 232 cells on 3 slices. Speed of nuclear translocations, straightness of cell body progression, cell body displacement per hour or percentage of time spent stationary were similar between both populations. Error bars are SEM. RMS: Rostral Migratory Stream; GCL: Granule Cell Layer; MCL: Mitral Cell Layer; GL: Glomerular Layer.

Thus, targeted electroporation of the lateral or the medial stem cell compartments labelled with high predominance the lineages that led to GCs and PGN, respectively. We used this model to study their migratory behaviour.

### Migratory behaviour of GC-P and PGN-P is indistinguishable

First, we compared GC-P and PGN-P during their tangential migration in the RMS using two-photon time-lapse imaging on acute brain slices at 5 days post lateral or medial electroporation (Fig. 1D, E). To increase the number of labelled cells for precise quantitative analyses, we used CRE-mRNA electroporation into R26 tdTomato transgenic mice. This approach allows specific targeting of the ventricular stem cell compartments at higher transfection efficiency (Bugeon et al., 2017).

Individual cells were manually tracked using Imaris software (Bitplane The instantaneous speed and other migratory parameters of the cells were then derived based on their tracked position at each frame of the time-lapse. Speed profiling of cell bodies (Fig. 1F), showed typical saltatory migration, alternating short stationary phases and fast nuclear translocations, for both populations (Movie 1). Speed of nuclear translocations, straightness of cell body progression, cell body displacement per hour and lengths of stationary phases were indistinguishable between both populations (Fig. 1G).

We then compared radial migration of both populations on acute brain slices at 7 dpe, a time point when large numbers of cells leave the RMS-OB and initiate radial migration to invade the OB target layers (Fig 2A, Movie 2). Quantification in the deeper half of the GCL during 5 hours showed that about 60% of the GC-P and 80% of PGN-P had migrated during the recording period (Fig. 2B, left). The difference in migration between the two populations was even larger when we analysed cells located in the superficial half of the GCL. Here, less than 30% of GC precursors, but still almost 80% of PGN-P, changed position during the recording period (Fig. 2B right). For the migratory fractions of both populations, radial migration was always saltatory, like in the RMS, but with lower speed of nuclear translocation as compared to tangential migration (Fig. 2C) (tangential: 75 μm/h; radial: 45 μm/h) (Fig. 2D). Comparison of the different migratory parameters at the population level showed no significant differences regarding the speed of nuclear translocation, displacement or percentage of time spent in stationary phases. Only a small but significant increase of straightness was detected, potentially meaning that GC-P have a slightly more exploratory behaviour. Next, we compared migratory parameters in the OB of GC-P and PGN-P at the cellular level. The example presented in Fig. 2E provides a high-resolution view of the sequence of events during which peaks in neurite length precede swelling formation and nuclear translocation (Fig. 2F). Neurite length and frequency of neurite extension were similar between populations. There was a tendency toward a decrease of swelling frequency and a small significant decrease in swelling duration between GC-P and PGN-P (Fig. 2G). This result may indicate a different swelling formation mechanism between these two populations.

**Figure 2:**
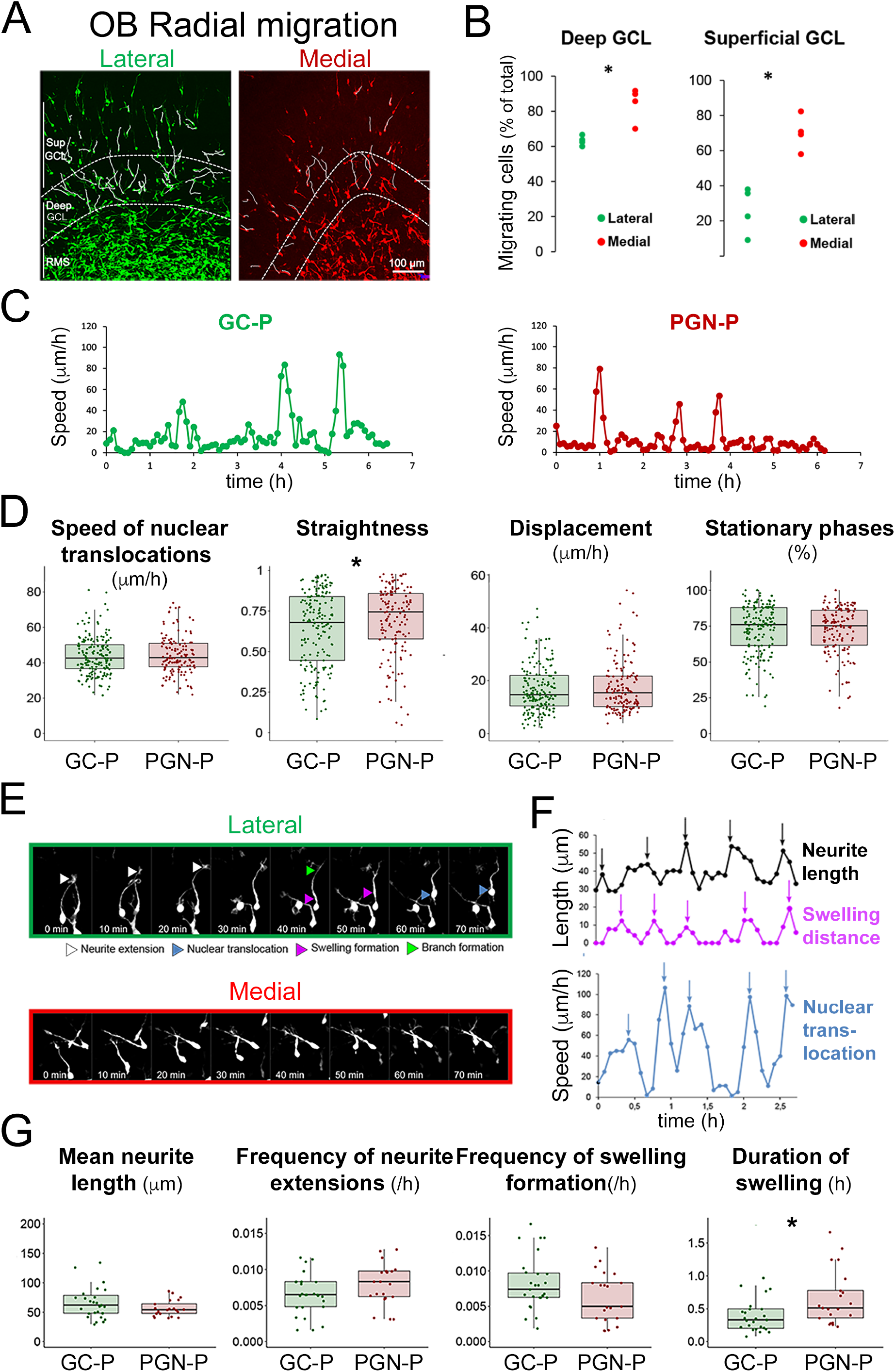
Characterization of radially migrating GC-P and PGN-P in the GCL. **A.** R26-Ai14 P0 animals received lateral or medial electroporation of a Cre encoding plasmid. The parameters of radial migration of TdTomato expressing neuroblasts in the GCL were then assessed on acute brain slices at 7 dpe. Example tracks (white lines) of radially migrating cells after lateral and medial electroporation. Dotted lines separate the RMS, deep GCL and superficial GCL. Scale bar: 100 μm. **B.** Percentage of cells migrating in the deep and superficial GCL during a 5-hour recording period. Cells with at least one period of active migration were considered. Each dot represents a slice. N = 4 slices, p<0.05; Wilcoxon rank sum test for means. **C.** Example of instantaneous speed recorded during a 6-hour movie for a lateral and a medial cell. **D.** Mean speed of nuclear translocations, straightness, displacement per hour and percentage of stationary phases for tangentially migrating lateral and medial cells. We found no significant differences regarding speed of nuclear translocation, straightness, mean displacement or percentage of time spent in stationary phases between both populations. N = 138 cells on 4 slices. Error bars are SEM. **E.** Snapshots of time-lapse two-photon imaging of radially migrating lateral (higher panel) and medial (lower panel) cells at high magnification. White arrowheads indicate the time of neurite extension, blue arrowheads indicate the occurrence of nuclear translocation, purple arrowheads indicate swelling formations and green arrowhead indicate a neurite branching appearance. **F.** Example showing the sequence of neurite extension, swelling formation and nuclear translocation periods for one cell. Upper chart: Neurite length and swelling distance are represented over time in μm. Lower chart: instantaneous speed of the cells in μm/h. Arrows indicate the peaks of each events. **G.** Mean neurite extension length, frequency of neurite extension events, frequency and duration of swelling formations. We found no major differences in neurite length, frequency of neurite extension, frequency of swelling between GC-P and PGN-P. The duration of swelling was slightly increased for PGN-P. N = 21 cells, 3 slices. For duration of swelling, p < 0.05, Wilcoxon rank sum test for means. Individual points represent cells. Error bars are SEM.

Overall, we conclude that GC-P and PGN-P share unique modes of tangential and radial migration towards and inside the OB. The significantly lower mobility of GC-P in the GC likely reflects the fact that this population is rapidly recruited to the circuitry and stop migrating. In contrast, PGN-P do not stop migration as they traverse the GCL and integrate the PGL. A clear morpho-kinetic predictor of neuronal fate during tangential or radial migration was not detected.

### Neuronal activity increases in GC-P during recruitment in the OB

To investigate the impact of neuronal activity on migration and recruitment of GC-P and PGN-P in the OB neurogenic system, we quantified their spontaneous calcium activity. CRE-mRNA was targeted to either the lateral or the medial stem cell compartments of R26 tdTomato/GCaMP6s mice. Acute slices were prepared and imaged between 6 and 10 dpe by 2-photon microscopy (Fig. 3A, B). Neuronal activity in GC-P and PGN-P was observed in three defined regions (Fig. 3C): first, in the RMS ‘elbow’ during tangential migration, before precursors enter the core of the OB; second, in the RMS-OB where the switch between tangential and radial migration occurs; third, in the deep GCL during radial migration.

**Figure 3:**
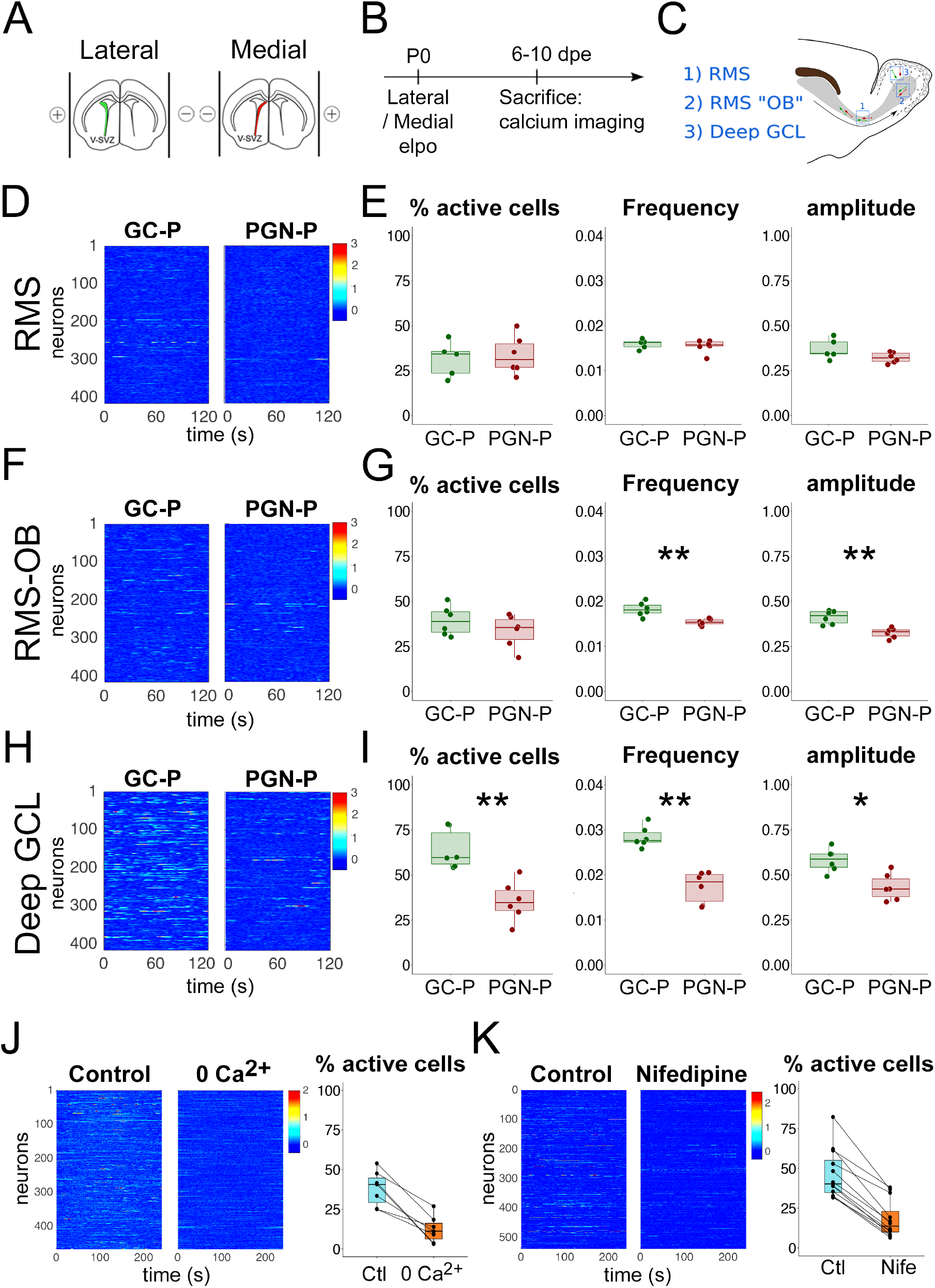
Spontaneous neuronal calcium activity in GC-P and PGN-P in the RMS, RMS-OB and deep GCL. **A.** Schematic representation of an olfactory bulb sagittal slice. **B.** Ai14/GCaMP6s P0 animals received lateral or medial electroporation of a Cre mRNAs. **C.** Calcium activity was recorded on acute brain slices between 6 and 10 dpe. Red and green cells represent medial and lateral cells migrating in the RMS, RMS-OB or in the deep GCL. **D.F.H.** Raster plot of calcium activity of GC-P (left) and PGN-P (right) recorded in the RMS (D), RMS OB (F) and deep GCL (H). **E.G.I.** Percentage of active cells, frequency and mean amplitude for GC-P and PGN-P in the RMS (E), RMS-OB (G) and in the deep GCL (I). Cells were defined as active if they showed at least one detectable calcium transient. The calcium activity of GC-P greatly increased upon arrival in the RMS-OB and the deep GCL, in contrast with PGN-P activity. N = 6 slices per condition. Error bars are SEM. **J.** Left: Raster plot of calcium activity of GC-P in the deep GCL in control and calcium-free medium (No Calcium + 2mM EGTA). Right: The percentage of active cells is highly decreased in calcium free medium. **K.** Left: Raster plot of calcium activity of GC-P in the deep GCL in control condition and nifedipine (10 μM) supplemented medium. Right: The percentage of active after incubation with the L-Type Voltage Gated Calcium Channels antagonist Nifedipine at 10 μM is highly decreased.

In the RMS, about one third of the imaged cells were active during a 2 minutes recording (i.e. at least one detectable transient) with no significant differences between medial and lateral slices (Fig. 3E). Similarly, the amplitude and frequency of calcium transients were comparable for the two populations, with a mean of 35% dF/F0 at a frequency of 0.016Hz (Fig. 3E). We then recorded spontaneous calcium activity in precursors located in the RMS-OB (6 slices from 3 animals, Fig. 3F). While the percentage of active cells was similar between GC-P and PGN-P, we observed a significant increase of the frequency and amplitude of calcium transient in GC-P. When precursors migrated in the deep GCL (Fig. 3 H), we observed an important increase of activity in GC-P compared to PGN-P: Indeed, the percentage of active cells was almost twice as large in GC-P, while there was respectively a 65% and 37% increase in amplitude and frequency of calcium transients between in GC-P compared to PGN-P.

We aimed at identifying the neurophysiological mechanisms underlying this increased activity of GC-P in the deep GCL. We used a wide spectrum of pharmacological interventions including the NMDA receptors antagonist D-APV, the AMPA receptors antagonist NBQX, the metabotropic glutamate receptor 5 antagonist MPEP, the GABAA receptors antagonist bicuculline, the glycine receptor antagonist strychnine (Fig 3-1). None of these antagonists significantly modified the percentage of active cells, or decreased the amplitude or frequency of calcium transients. However, perfusion of a calcium-free medium on OB slices led to a strong decrease in the percentage of active cells (41 vs 11 %, p=0.004, Fig. 3J, Fig 3-1A) suggesting that the source of calcium was mainly extracellular. Moreover, blocking specifically L-type voltage gated calcium channels with nifedipine also abolished most of calcium activity (Fig. 3K, Fig 3-1 B). Taken together, these two results suggest that the calcium activity increase seen in GC-P was mainly due to a depolarization, leading to the opening of voltage dependent L-type calcium channels and an entry of extracellular calcium.

**Figure 3-1:**
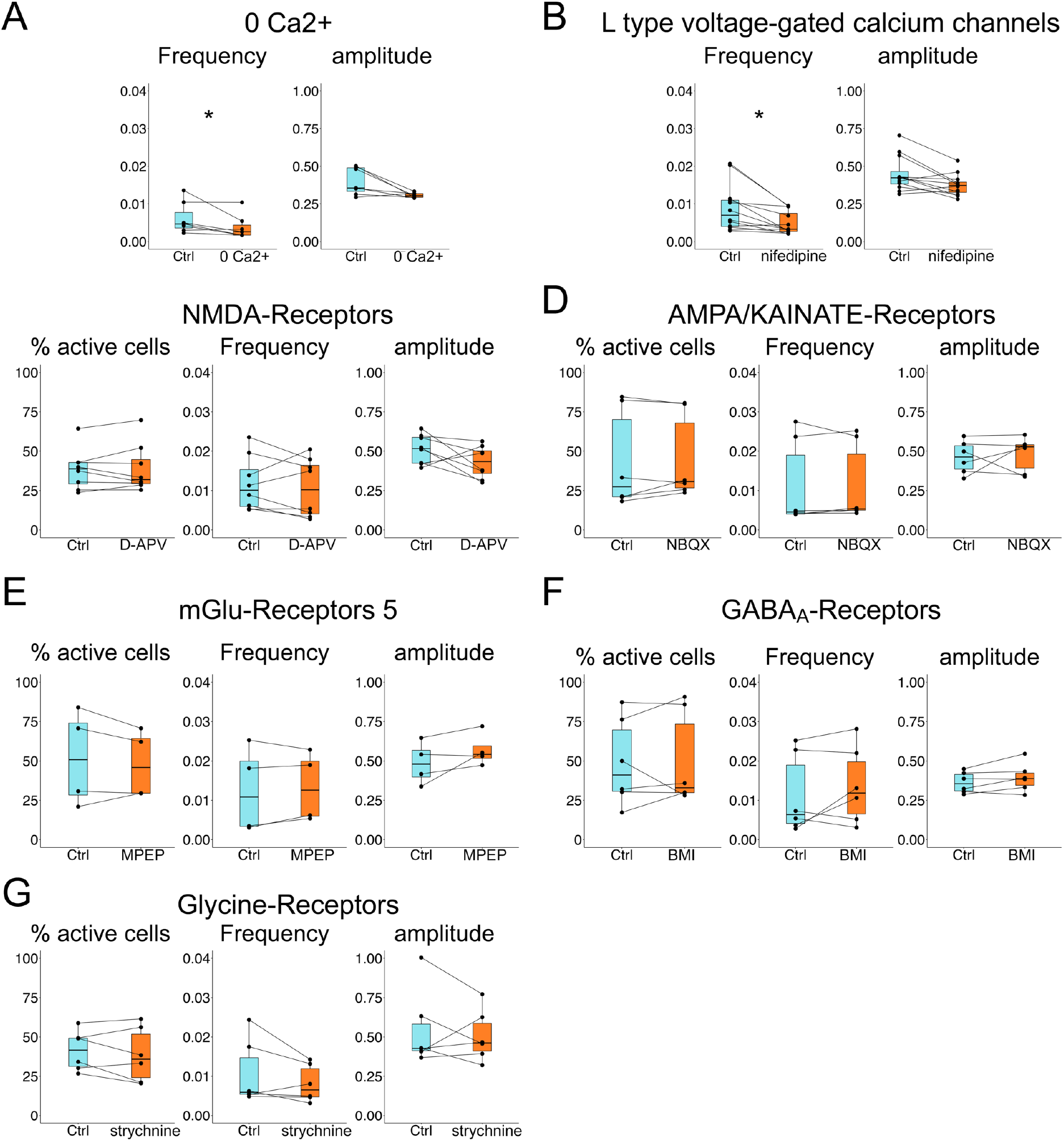
Spontaneous neuronal calcium activity in GC-P in the deep GCL is not affected by classical antagonists. Incubation with a calcium free medium decrease the frequency of calcium transients. (A) The L-type voltage-dependent calcium channel blocker nifedipine decrease the frequency of calcium transients. (B) The NMDA receptors antagonist D-APV (C), AMPA/Kainate receptors antagonist NBQX (D), metabotropic glutamate receptor 5 antagonist MPEP (E), GABAA receptors antagonist Bicuculline (F) and glycine receptor antagonist strychnine (G) do no modify the percentage of active cells or the amplitude of calcium transient.

Thus, after entering the RMS in the OB, and especially when they start exiting the RMS, GC-P, but not PGN-P, showed an L-type calcium channel dependent increase in neuronal activity. These results led us to hypothesize that the depolarization of GC-P during the transition from the RMS into the GCL is critical for their recruitment.

### Neuronal activity is needed for recruitment and survival of GC-P

To test if neuronal activity controls the recruitment of precursors from the RMS to the OB layers, we interfered with the activity of arriving GC-P. It was demonstrated that overexpression of the inward rectifying potassium channel Kir2.1 could highly affect neuronal activity by lowering the resting membrane potential, therefore altering neuronal excitability (De Marco Garcia et al., 2011; Xue et al., 2014). Over-expression of Kir2.1 in GC-P was performed by electroporation of a Kir2.1 expression plasmid into the lateral ventricular wall (Fig 4A).

**Figure 4:**
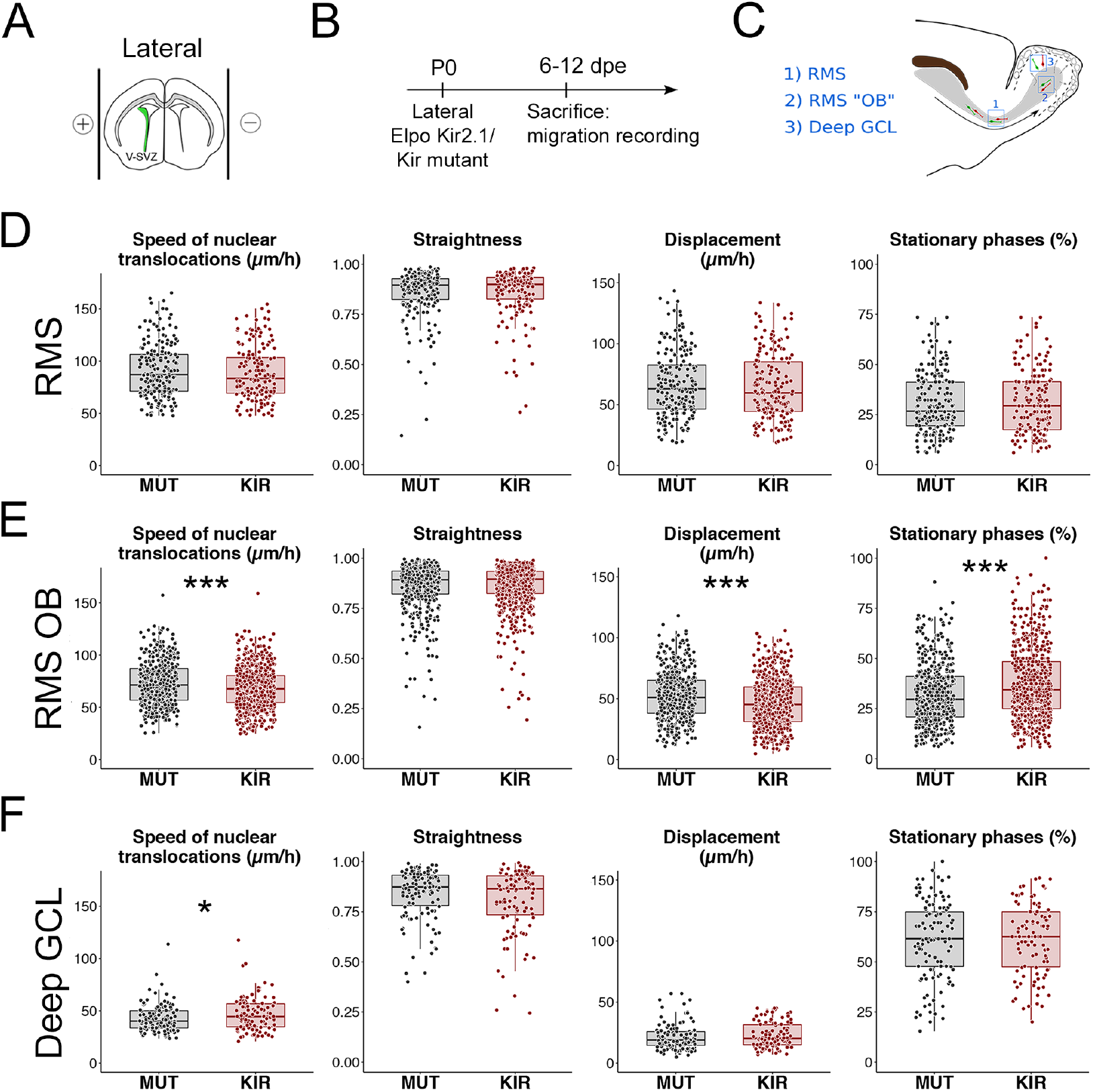
Effect of Kir2.1 expression on GC-P migration. **A.** P0 CD1 animals received lateral electroporation with either a Kir2.1 mutant or a functional Kir2.1 encoding plasmid. **B.** Mice were then sacrificed between 6 to 12 dpe. **C.** the parameters of migration of GC-P were then assessed on acute brain slices between 6 and 12 dpe in the RMS, RMSOB and deep GCL. **D.E.F.** Quantification of the speed of nuclear translocation, the straightness, the displacement and the percentage of cells in stationary phase in the RMS (D), RMS-OB(E) and in the deep GCL (F). In the RMS-OB, when Kir2.1 is expressed, migration is affected: the speed of nuclear translocation and the displacement are decreased and the percentage of stationary cells is increased. A slight increase of the speed of nuclear translocation is observed in the deep GCL

First, we investigated if blocking neuronal activity of GC-P affects their migration. In the RMS (Fig. 4C), a region where spontaneous neuronal activity in GC-P was low and indistinguishable from PGN-P (Fig. 3D), expression of Kir2.1 had no effect on the speed of nuclear translocations, straightness, cell displacement and stationary phases (Fig. 4D). Next, we analysed migration in the RMS-OB, a region where neuronal activity is increased in GC-P (Fig. 3G). In this region, expression of Kir2.1 led to a decrease in the speed of nuclear translocation, the displacement per hour and to an increase in the percentage of cells in stationary phase. Finally, we analysed GC-P in the deep GCL. Here, Kir2.1 expression had again no detectable effect on straightness, displacement and stationary phases (Fig. 4F). Only the speed of nuclear translocations showed a small but significant increase (Fig. 4F).

We then tested if neuronal activity controls the recruitment of GC-P and PGN-P in the OB layers. To test this hypothesis, we overexpressed Kir2.1 in GC-P and PGN-P by lateral or medial electroporation and analysed neuronal distribution in the OB at 8 and 12 dpe, when migratory precursors leave the RMS-OB and settle in the OB (Fig. 5A-B). At 8 dpe, the number of Kir2.1 expressing cells was significantly higher in the RMS-OB whereas less cells were present in the GCL (Fig. 5C). This effect was specific to GC-P, as distribution was unaffected in PGN-P after medial electroporation (Fig. 5D). At 12dpe, when more cells entered the OB, this altered distribution in the GC population was slightly more pronounced (Fig. 5c) and, again, not detectable in PGN-P (Fig. 5D). At both time points cell distribution in the GL was unaffected (Fig. 5CD).

**Figure 5:**
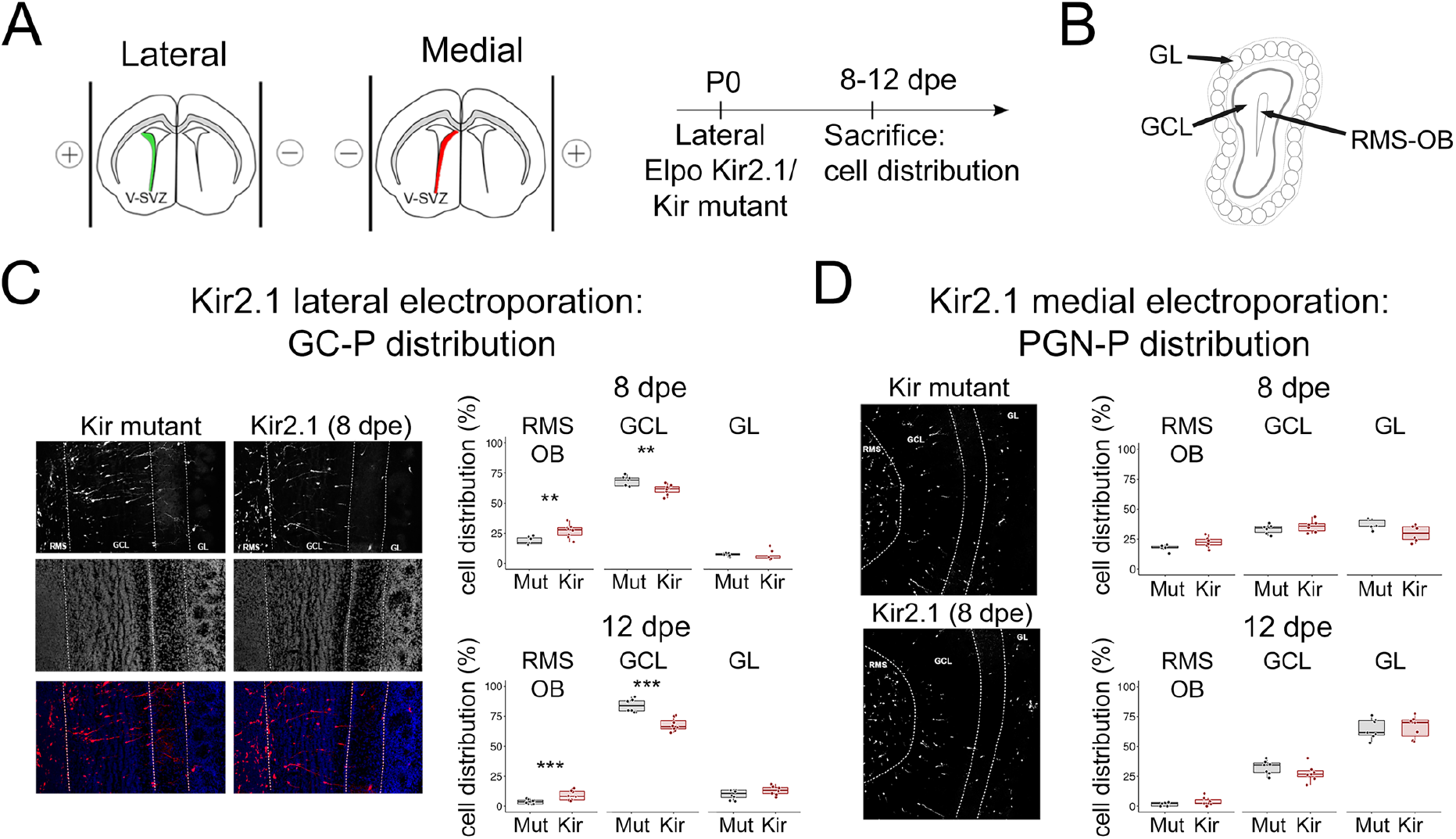
Effect of Kir2.1 expression on cellular distribution. **A.** P0 CD1 animals received lateral or medial electroporation with either a Kir2.1 mutant or a functional Kir2.1 encoding plasmid. Mice were then sacrificed between at 8 or 12 dpe. and the cellular repartition of TdTomato cells was assessed. **B.** The cellular distribution was observed on fixed coronal brain slices and cells attributed to the RMS, GCL or GL. Red and green cells represent medial and lateral cells migrating in the RMS, RMS-OB or in the deep GCL. **C.** Left: OB coronal sections of Kir2.1 mutant (Mut) and Kir2.1 (Kir) laterally electroporated animals at 12dpe. Dotted lines separate OB layers. DAPI stains cell nuclei. Scale bar: 500 μm. Right: Quantification of the cellular repartition at 8 dpe and 12 dpe in each OB layer showing an increased percentage of Kir2.1 cells in the RMS and a decrease in the GCL. **D.** Left: OB coronal sections of KIR2.1 mutant and Kir2.1 medially electroporated animals at 12dpe. Dotted lines separate OB layers. Scale bar: 500 μm. Right: Quantification of the cellular repartition at 8 dpe and 12 dpe in each OB layer. No significant effect on cellular distribution was observed in this condition.

Finally, we asked if blocking activity and/or the subsequent misdistribution of GCs had an impact on cell survival. To this end we over-expressed Kir2.1 and analysed the expression of cleaved caspase 3 (c-caspase 3), a marker of apoptosis at 8 and 12 dpe in the RMS, the GCL and the GL (Fig. 6AB). In agreement with previous studies, we found that overall level of c-caspase 3 expressing cells was very low (Petreanu and Alvarez-Buylla, 2002; Yamaguchi and Mori, 2005; Platel et al., 2010). However, 8 days after lateral electroporation, we detected a slight but significant increase of c-caspase 3 immunoreactive cells among GC-P expressing Kir2.1 in the GCL. At 12 dpe, this effect was even more pronounced in the GCL (Fig. 6C) and we observed a strong but not significant tendency in the RMS-OB. Importantly, we did not observe any increase in apoptosis after expression of Kir2.1 in PGN-P at 8 or 12 dpe (Fig. 6D).

**Figure 6:**
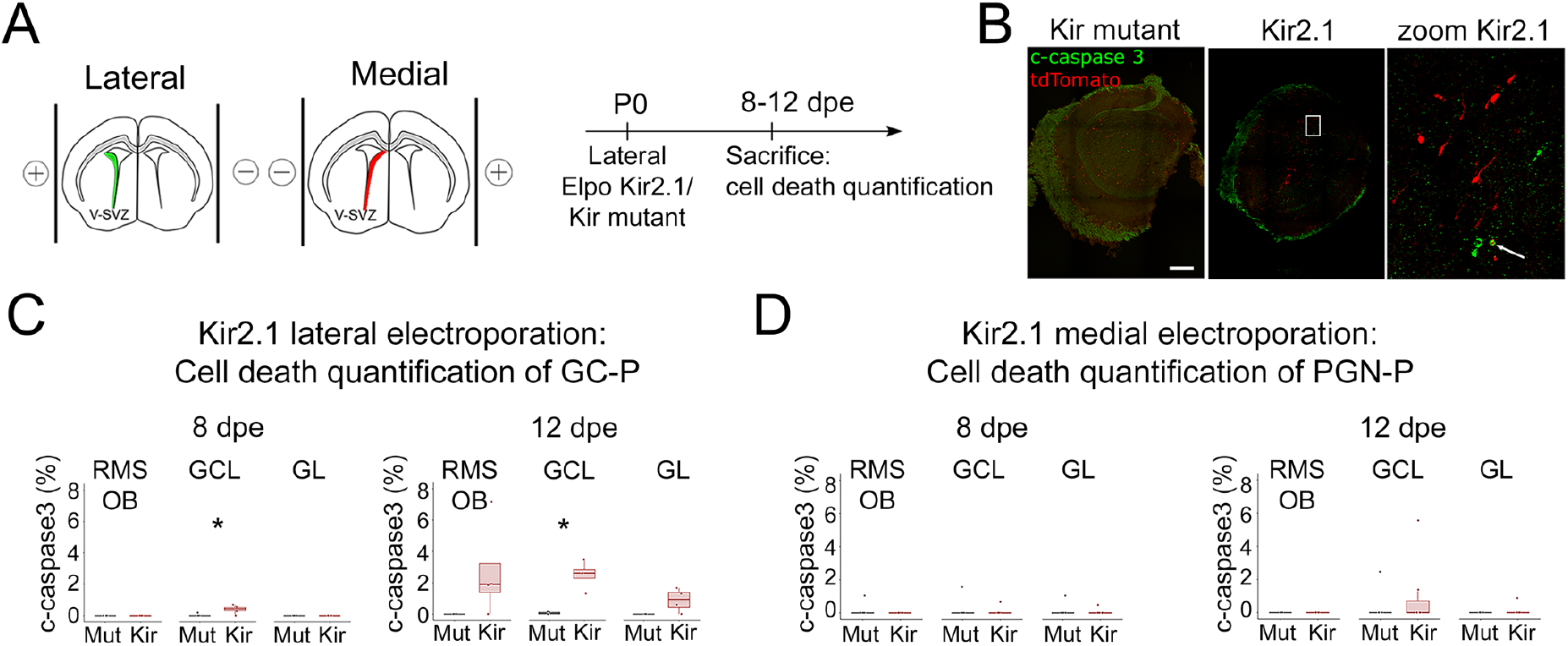
Effect of Kir2.1 expression on neuronal cell death. **A.** P0 CD1 animals received lateral or medial electroporation with either a Kir2.1 mutant or a functional Kir2.1 encoding plasmid. Mice were then sacrificed at 8 and 12 dpe and the level of apoptotic cell death in TdTomato cells quantified. **B.** OB coronal sections of Kir mutant and Kir2.1 labeled with cleaved caspase 3 at 12dpe. Note the presence of a double labeled cell in the Kir condition **C.D.** Quantification of the percentage of cleaved caspase 3 positive cells among lateral (C) or medial (D) electroporation of Kir2.1 (Kir) or Kir2.1 mutant (Mut) cells per OB region at 8dpe or 12 dpe. There is an increase of cleaved caspase 3 positive cells after lateral electroporation of Kir2.1 in the GCL at 8 dpe and 12 dpe but not after medial electroporation.

Overall, we conclude that activity in GC-P is important for migration in the RMS-OB, positioning in the GCL and survival while PGN-P are not affected by the inhibition of activity. Taken together, these results suggest that activity in GC-P at the exit of the RMS is a controlling factor in the recruitment and positioning process in the OB.

## Discussion

Combining targeted electroporation and time-lapse two-photon imaging, we were able for the first time to compare and analyse migrating precursors of the two main OB interneuron subtypes. Our deep characterization of the tangential and radial migration of these two cell types did not show any obvious morphologic or kinetic parameter that allows to differentiate them, suggesting that GC and PGN-P share the same mode of migration to reach the olfactory bulb. However, we identified a dramatic increase of calcium activity in GC-P at the time of switching between tangential to radial migration. This calcium activity was found to be mostly due to a voltage-dependent extracellular calcium entry through L-type calcium channels meaning that this precursors were depolarised by an unknown factor. Blocking this activity by dampening neuronal excitability altered GC-P migratory speed in the RMS OB, which is the place where tangential to radial migration switch occurs. Interestingly, blocking activity also led to a progressive accumulation of GC-P in the RMS OB at the expanse of the GCL and to a non-negligible increase in cell death in radially migrating GC-P. In contrast, blocking neuronal activity in PGN precursors did not affect their cellular distribution or cell death during migration. Overall, these experiments suggest that neuronal activity in GC precursors is necessary for the proper switching to radial migration, while this is not the case for PGN-P.

It was shown in the OB that olfactory training (Mouret et al., 2008) or overexpression of NaChBac (Lin et al., 2010) could increase recruitment during synaptic integration. In sharp contrast, it was recently demonstrated that in physiological condition selection during synaptic integration is nearly absent (Platel et al., 2019) pointing to a first selection step occuring before integration (Biebl et al., 2000; Platel et al., 2010). In agreement with these latter studies, our results suggest that survival of neuroblasts during radial migration is dependent on neuronal activity. Importantly, this phenomenon seems to be subpopulation specific. While it is known that layer positioning of different cortical interneurons depends on neuronal activity (De Marco Garcia et al., 2011) (Lodato et al., 2011), it is unclear to which extent this is a result of altered migration or differential cell death.

Our study also show a subtype-specific effect of neuronal activity during the switch between tangential to radial migration. However, we could not identify a specific signal explaining the large increase in neuronal activity at the RMS exit for GC-P. It was found for migrating glutamatergic cortical neurons that transient synaptic connections occur during migration and that these connections were impacting migration (Ohtaka-Maruyama et al., 2018). In our pharmacological screen, neither glutamate nor GABA nor glycine receptors’ antagonists were able to block the large rise in calcium activity. A tempting hypothesis would be that rather than being dependent on extracellular signals (such as neurotransmitter), this activity could be the consequence of neuroblast maturation. Interestingly, our observations suggest that cells can migrate for a significant time (at least 8 hours) in the RMS of the OB before exiting in the granule cell layer. In this hypothesis, migrating cells in the RMS would need to reach a certain threshold of maturation, allowing proper intrinsic activity, which would in turn favor their switch to radial migration.

Contrary to GC precursors, PGN precursors were not affected by Kir2.1 mediated hyperpolarization suggesting that the recruitment of future PGNs is not activity-dependent. This is in line with findings that knocking out serotonin receptors did no affect cell density in the glomerular layer (Garcia-Gonzalez et al., 2017) and results showing that the migration of PGN precursors in the GL does not depend on sensory activity (Liang et al., 2016). Of note, this later study also showed that PGN neurons migrate for an extended period in the GL while most GCs had settled. This is coherent with the hypothesis that the maturation of GCs and PGNs during migration follows a different time course.

To our knowledge there is no candidate factor known to regulate the addressing of PGN to their specific layer. New advances in single cell transcriptomics coupled to our specific targeting of the two subpopulations might allow to decipher which molecular pathways differ between PGN and GC precursors during their radial migration.

In conclusion, we show that intrinsic neuronal activity early during neuronal development, when neurons are still migrating, can control the recruitment of specific interneuron subtypes to their target layer.

## Methods

### Animals

All mice were treated according to protocols approved by the French Ethical Committee (#5223–2016042717181477 v2). Mice were group housed in regular cages under standard conditions, with up to five mice per cage on a 12 hr light–dark cycle. CD1 (Charles-River, Lyon, France), Ai14 (Rosa TdTomato) transgenic reporter (Jackson Laboratories, stocknumber 007914) and GCaMP6s (Jackson Laboratories, stock-number 028866) mice were used. All experiments were performed on males and females.

### Plasmids and mRNA preparation

pCAG-Kir2.1-T2A-tdTomato (https://www.addgene.org/60598/) (Xue et al., 2014), pCAG-Kir2.1Mut-T2A-tdTomato (https://www.addgene.org/60644/). The pCAGGS-EGFP vectors was derived from pCX-MCS2 (Morin et al., 2007). All plasmids were used at a concentration of 5 μg/μl (0.1% Fast Green). mRNAs were provided by Miltenyi Biotec (Miltenyi Biotec): CRE Recombinase mRNA (130-101-113) (a kind gift from A. Bosio, Miltenyi Biotec). For electroporation, mRNAs were diluted in RNAse-free PBS at a concentration of 0.5 μg/μl (same range of molarity compared with 5 μg/μl DNA plasmid).

### Immunohistochemistry and image analysis

For histological analysis, pups were deeply xylazine/ketamine anaesthetized. Perfusion was performed intracardiacally with a solution of 4% paraformaldehyde in PBS. The brain was dissected out and incubated overnight in the same fixative at 4°C. Sections were cut at 50 μm using a microtome (Microm). Floating sections were first incubated overnight at 4°C with the following primary antibody: cleaved-Caspase3 antibody (cell signalling technology), before incubation 2 h at room temperature with the corresponding fluorescent labelled secondary antibody. Before mounting, cell nuclei were stained with Hoechst 33258. Optical images were taken either using a fluorescence microscope (Axioplan2, ApoTome system, Zeiss) or a laser confocal scanning microscope (LSM510 or LSM780, Zeiss).

For quantifications of cell repartition, layers were manually drawned based on the nuclear staining. Then, the CellCounter plugin from ImageJ was used to manually count cells in each layer. All experiments and quantifications were performed blindly to experimental groups.

### Postnatal electroporation

Postnatal electroporation was performed as described previously (Boutin et al., 2008; Bugeon et al., 2017). Briefly, P0-P1 pups were anesthetized by hypothermia. A glass micropipette was inserted into the lateral ventricle, and 2 μl of plasmid or RNA solution was injected by expiratory pressure using an aspirator tube assembly (Drummond). Successfully injected animals were subjected to five 95 V electrical pulses (50 ms, separated by 950 ms intervals) using the CUY21 edit device (Nepagene, Chiba, Japan) and 10 mm tweezer electrodes (CUY650P10, Nepagene) coated with conductive gel (Control Graphique Medical, France). Electrodes were oriented according to the desired targeting: for an electroporation of the left hemisphere, the cathode was placed on the left side and anode on the right side for lateral electroporation; for medial electroporation cathode on the right side and anode on the left side. Due to the lower number of cells generated from the medial V-SVZ (Ventricular-SubVentricular Zone), electroporations were preferentially performed at P0, when it is the most efficient. Electroporated animals were then reanimated in a 37°C incubator before returning to the mother.

### Acute brain slices

Seven days after electroporation, P7-8 animals were ketamine/xylazine anesthetized, and perfused intracardiacally with cold (4°C), oxygenated (95% O2 / 5% CO2) dissection solution (250mM Sucrose, 3 mM KCl, 1.25 mM NaH2PO4, 3 mM MgSO4, 10 mM D-Glucose, 26 mM NaHCO3 and 0.5 mM CaCl2). The brain was then quickly removed, and glued on a vibratome platform. 300 μm thick sagittal slices were taken with a vibratome (Leica VT1200S) in chilled (4°C), oxygenated (95% O2 / 5% CO2) dissection solution. Thick sections were then placed in oxygenated DMEM, high glucose, GlutaMAX (Gibco) or ACSF (Artificial CerebroSpinal Fluid) (124mM NaCl, 3mM KCl, 1.25 mM NaH2PO4, 1.3 mM MgSO4, 10mM D-Glucose, 26 mM NaHCO3 and 2mM CaCl2) at room temperature for at least 1 hour prior to imaging.

### Two-photon time-lapse imaging

Acute brain slices were imaged with a two-photon ZEISS 7MP, equipped with a 20X objective (1,0 NA) and a Mai-tai laser (ONE BOX TI:SAPPHIRE LASERS, Spectra-Physics). Slices were imaged in a flow-through warming chamber (Warner instruments, Open Diamond Bath Imaging Chambers, RC-26G) and continuously superfused (Compact Peristaltic Pump, Harvard Apparatus; ~1 ml/min) with the appropriate recording solution bubbled with 95% O2/5% CO2 using a dual automatic temperature controller with an in-line heater (Warner Instruments). Slices were maintained with a slice anchor (Warner instruments, SHD-26GH/10).

For migration, z-stacks (step size: 3μm) were taken every 2 minutes for tangential migration and every 5 minutes for radial migration. All images were 1024X1024 pixels large, with an axial resolution of 0.52 μm. Z-stack maximum projections were then realized on every time points, and registered for horizontal drifting using the StackReg plugin from Image J software.

For calcium imaging, single plane time-lapse were acquired every 1 s, for at least 2 minutes. Only movies with low Z drifting were analyzed. Again, movies were registered using the StackReg plugin from ImageJ.

All acquisitions were done at least 50 μm below the surface of the slice, to avoid strong slicing artefacts.

### Cell tracking and migration analysis

Individual cells were manually tracked using Imaris software (Bitplane). Cell positions were then saved in individual excel files for each movie. The parameters of saltatory migration were then analyzed in R.

Instantaneous speed at time t was calculated by dividing the distance separating the positions at t-1 and t+1 by two time intervals. A cell was considered as moving when this speed reached 20 μm/h corresponding to a displacement of 1.3 μm per time frame (for a 5 minutes time interval), a distance sufficient to avoid noisy movements detection. From the speed vector, we could determine phases of active migration and phases of relative immobility (stationary phases). This allowed the calculation of the mean speed of nuclear translocations corresponding to the mean of all speed above 20 μm/h and the percentage of stationary phases which is the percentage of time points with a speed lesser than 20 μm/h. These parameters were calculated from the first migratory phase to the last one. Displacement per hour was calculated by dividing the total distance covered during the recording divided by the time required to travel this distance. Straightness was defined as the ratio between the displacement vector length and the cumulative distance traveled. Cells with no migratory phase were not taken into account. Cells with a single migratory phase were excluded from the calculation of the percentage of stationary phases.

For quantifying the percentage of cells migrating in the superficial vs deep granule cell layers, we considered croped our time lapses make them all 5 hours long. Then the GCL was divided in two equal halves (deep and superficial). A cell was considered migrating if it showed a least one migratory period during the 5 hours of the time lapse.

Neurite length and swelling formation distance were measured manually with ImageJ. Mean neurite length extension was calculated over periods when neurite length was above the mean neurite length plus two standard deviation (SD). Similarly, the frequency of neurite extensions was calculated as the number of neurite extensions (mean neurite length+2SD) per hour. The threshold for the swelling distance was set at 4 μm, above this value it was considered as a swelling formation. From this, the frequency of swelling formation was defined as the number of swelling formation per hour. The mean duration of swelling was the mean time spent with a swelling formation.

### Calcium imaging analysis

All calcium imaging experiments were performed on GCaMP6s/Ai14 mice electroporated with the RNA Cre recombinase. The expression of this calcium sensitive probe in electroporated cells allows the visualization of intracellular calcium fluctuations.

Calcium time-lapse movies were averaged over time to obtain cell contours. Averaged images were then segmented to obtain a binary mask that was treated with the “Analyze Particles” methods in ImageJ software. Automatically drawn ROIs (Region Of Interest) were then manually curated to remove lost cells, duplicates or mistakes. The mean intensity of green fluorescence for each ROI was measured over time and saved as text file. A homemade Matlab interface was used to import, filter (with running average) and normalize intensities. Calcium transients were automatically detected by the software, with a fixed threshold of 25% dF/F0. The end of each calcium transients was defined as more than 3 time points with a consecutive decrease of amplitude. Each peak was then manually curated to avoid systematic errors from the software. The mean amplitude and frequency of calcium transients from each cell were then calculated and saved.

### Pharmacology

Antagonists were bath applied at the following concentrations: D-APV (Tocris Bioscience, 50μM), NBQX (Tocris Bioscience, 10μM), MPEP (Tocris Bioscience, 30μM), Bicuculline (Tocris Bioscience, 10μM), Strychnine (Tocris Bioscience, 20μM), Nifedipine (Tocris Bioscience, 10μM).

Calcium-free solution was prepared from modified ACSF (124mM NaCl, 3mM KCl, 1.25 mM NaH2PO4, 1.3 mM MgSO4, 10mM D-Glucose, 26 mM NaHCO3, and 1 mM EGTA).

### Statistics

Statistical analyses were performed using R software and R Commander Package (https://CRAN.R-project.org/package=Rcmdr). Data are presented as boxplots using the ggplot2 package.

Non-parametrical Wilcoxon rank sum test was used to assess mean differences between two groups.. Levene’s test was used to assess equality of variances which were considered statistically different if p<0.05. For figure 3 and figure 3-1 we used a paired Wilcoxon test. Probability assignment: p > 0.05 (not significant, ns), 0.01< p < 0.05 (*), 0.001< p < 0.01 (**) and p < 0.001 (***).

## Acknowledgements

The authors thank the Cremer lab, Jenelle Wallace and Dominique Debanne for critical reading of the manuscript. We are particularly grateful to the local PiCSL-FBI core facility (IBDM, AMU-Marseille) supported by the French National Research Agency through the « Investments for the Future’ program (France-BioImaging, ANR-10-INBS-04) as well as the IBDM animal facilities.

This work was supported by Fédération pour la Recherche sur le Cerveau (FRC) to HC, Agence National pour la Recherche (grant ANR-13-BSV4-0013), Fondation pour la Recherche Medicale (FRM) grants ING20150532361, to HC and FDT20160435597 to SB, Fondation de France (FDF) grant FDF70959 to HC.

